# NF-κB dependent gene expression and plasma IL-1β, TNFα and GCSF drive transcriptomic diversity and CD4:CD8 ratio in people with HIV on ART

**DOI:** 10.1101/2025.02.14.638232

**Authors:** Yingfan Wang, German G. Gornalusse, David A. Siegel, Alton Barbehenn, Rebecca Hoh, Jeffrey Martin, Frederick Hecht, Christopher Pilcher, Lesia Semenova, David M Murdoch, David M Margolis, Claire N. Levy, Keith R. Jerome, Cynthia D Rudin, Florian Hladik, Steven G. Deeks, Sulggi A. Lee, Edward P Browne

## Abstract

Despite antiretroviral therapy (ART), people with HIV (PWH) on ART experience higher rates of morbidity and mortality vs. age-matched HIV negative controls, which may be driven by chronic inflammation due to persistent virus. We performed bulk RNA sequencing (RNA-seq) on peripheral CD4+ T cells, as well as quantified plasma immune marker levels from 154 PWH on ART to identify host immune signatures associated with immune recovery (CD4:CD8) and HIV persistence (cell-associated HIV DNA and RNA). Using a novel dimension reduction tool - Pairwise Controlled Manifold Approximation (PaCMAP), we defined three distinct participant transcriptomic clusters. We found that these three clusters were largely defined by differential expression of genes regulated by the transcription factor NF-κB. While clustering was not associated with HIV reservoir size, we observed an association with CD4:CD8 ratio, a marker of immune recovery and prognostic factor for mortality in PWH on ART. Furthermore, distinct patterns of plasma IL-1β, TNF-α and GCSF were also strongly associated with the clusters, suggesting that these immune markers play a key role in CD4+ T cell transcriptomic diversity and immune recovery in PWH on ART. These findings reveal novel subgroups of PWH on ART with distinct immunological characteristics, and define a transcriptional signature associated with clinically significant immune parameters for PWH. A deeper understanding of these subgroups could advance clinical strategies to treat HIV-associated immune dysfunction.

## Introduction

Despite effective antiretroviral therapy (ART), people with HIV (PWH) on ART experience higher rates of morbidity and mortality compared to age-matched HIV negative controls [1–3]. PWH on ART also exhibit persistent changes to the immune system that do not fully normalize despite years of therapy, including a reduced CD4:CD8 T cell ratio, elevated levels of inflammatory immune markers, reduced levels of naïve T cells and elevated proportions of activated and exhausted T cells [2,4–7]. The mechanisms that drive ongoing immune dysfunction during ART are not well understood but may represent the sequelae of permanent damage caused to the immune homeostasis during untreated infection, ongoing stimulation of the immune system by viral RNA or antigen expression, or toxicity from ART. Previous studies have also demonstrated that the CD4:CD8 ratio is predictive of non-AIDS morbidity in PWH on ART, highlighting the clinical significance of this parameter [8–10].

Understanding the immune phenotypes of PWH and how these phenotypes are correlated with clinical comorbidities and HIV persistence will help to clarify the mechanisms of persistent immune dysfunction and potentially inform novel HIV cure strategies. In this study, we examine a published multimodal dataset derived from a cohort of 154 PWH along with additional unpublished data [11]. This dataset consists of transcriptomic profiles for CD4 T cells, total HIV DNA reservoir frequency, viral RNA expression, “intact” viral reservoir frequency, and plasma concentration levels for seven markers (IL-10, IL-1β, Pentraxin, TNF-α, IP-10, G-CSF, and sTLR4). From this dataset, we use a novel dimension reduction tool, Pairwise Controlled Manifold Approximation (PaCMAP), to identify three clusters of PWH with distinct transcriptomic profiles.

Furthermore, this clustering was associated with different levels of expression for NF-κB response genes in the CD4 T cells, plasma IL-1β, GCSF and TNF-α levels, and with the CD4:CD8 T cell ratio. These data indicate that virological or immunological events prior to ART establish a persistent dynamic of altered cytokine secretion that leads to divergent levels of chronic NF-κB signaling and transcriptomic perturbations in PWH. These processes may be important for understanding the persistence of the HIV reservoir and disruption of immune homeostasis in PWH on ART.

## Results

### Dimension reduction identifies three transcriptomic clusters within a cohort of ART-treated PWH

To define distinct immune phenotypes of people with HIV, we analyzed bulk RNA sequencing data from peripheral CD4+ T cells, plasma immune marker data, and HIV reservoir data, from 154 PWH on ART from the UCSF SCOPE and Options cohorts [11]. As previously described, the study included mostly male (95%) PWH who were virally suppressed on ART for a median of 4.9 years, consistent with our San Francisco-based population of PWH (**Table 1**, **Table S1**). Plasma levels of seven immune markers (IP-10, G-CSF, Pentraxin-3, IL-1β, IL-10, TNF-α and sTLR4) were measured, and the transcriptome of total CD4+ T cells for each participant was analyzed by bulk RNAseq (**Figure 1A**). To first analyze the overall structure of the dataset, and to potentially find distinct clusters of PWH, we applied a recently developed dimension reduction tool - Pairwise Controlled Manifold Approximation (PaCMAP) to the dataset [12]. PaCMAP accurately preserves both local and global data structure in high dimensional datasets. We first performed feature selection, in which the top 2000 most variable genes were considered for used with PaCMAP. Interestingly, when we projected the transcriptomes of the CD4 T cells from each PWH across a two-dimensional space, we observed three distinct clusters of PWH (**Figure 1B**). Notably, other dimension reduction methods such as Principal Component Analysis (PCA), T-Distributed Neighbor embedding (tSNE) and Uniform Manifold Approximation and Projection (UMAP) did not display clear global structure that matched PaCMAP, although PWH from each of the PaCMAP clusters occupied mostly non-overlapping regions in these projections (**Figure S1**). The PaCMAP clusters were robust across multiple random starting states (**Figure S2**) and consisted of 94 PWH (cluster 1), 51 PWH (cluster 2), and 9 PWH (cluster 3) (**Figure 1C**). These analyses suggest that there are three main subgroups of PWH within the cohort that have distinct transcriptomic phenotypes for their CD4 T cells, with most PWH being members of two larger clusters, and a smaller number being part of a third cluster.

**Figure 1:**
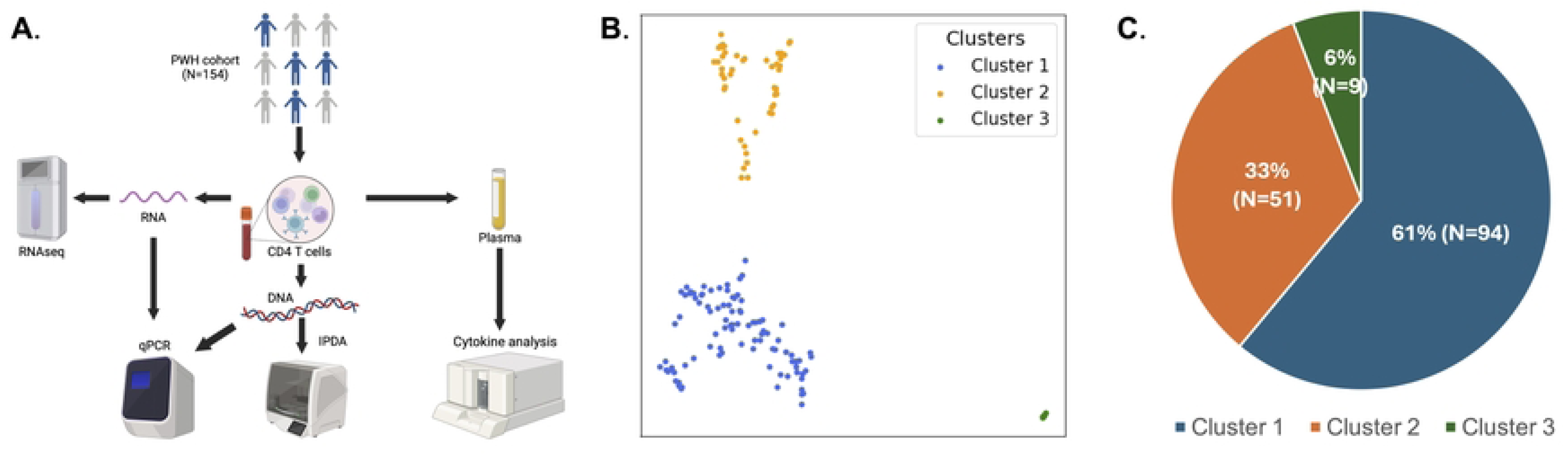
PaCMAP analysis of CD4 T cell transcriptomes identifies three clusters of people with HIV (PWH) on ART. **A.** Schematic overview of experimental design. **B.** CD4 T cell transcriptomes from a cohort of 154 people with HIV on ART were analyzed using Pairwise Controlled Manifold Approximation Projection (PaCMAP) [29]. The overall cluster structure is shown, with the three major clusters labeled (Cluster 1 -blue, Cluster 2 – orange, Cluster 3 - green). Each datapoint represents an individual PWH. **C.** Pie chart showing composition of the overall dataset with respect to cluster size. Percentage and number of PWH in each cluster is shown.

**Table 1:** Cohort Characteristics.

| Descriptive characteristic | Total (N=154) |
| --- | --- |
| Male (%) | 146 (95%) |
| Age (years) | 47 (25-70) |
| Years of therapy | 4.9 (0.6-16.6) |
| Nadir CD4+ T cell count (cells/mm<sup>3</sup>) | 336 (2-1500) |
| Maximum pre-ART HIV RNA (log<sub>10</sub>copies/mL) | 47260 (40-4494698) |
| HIV total DNA (log<sub>10</sub>copies/10<sup>6</sup> CD4+ T cells) | 13.3 (0-561.4) |
| HIV unspliced RNA (log<sub>10</sub>copies/10<sup>6</sup> CD4+ T cells) | 828.0 (0-16550.5) |

### Identification of differentially expressed genes that drive cluster membership

We next investigated the biological basis for the transcriptomic clusters. To compare the expression of cellular genes across the dataset, we examined pairwise differential gene expression between each of the clusters. Specifically, we used DESeq2 [13] to identify differentially expressed genes (DEGs) between each cluster combination then visualized the sets of DEGs as volcano plots (**Figure 2A, Table S2, Table S3, Table S4**). We identified 1377 genes that were differentially expressed (log_2_FC>1, Pval_adj_<0.05) between clusters 2 and 1, 5161 genes that were different between clusters 3 and 2, and 5815 genes that were different between clusters 3 and 1. In general we observed that more genes were upregulated in cluster 3 and cluster 2 compared to cluster 1, and that more genes were upregulated in cluster 3 compared to cluster 2. The volcano plots of differentially expressed genes between the clusters also indicated the polarized nature of the changes in gene expression (**Figure 2A**). We also visualized the expression level for a set of the most highly differentially expressed genes between the clusters as a heatmap (**Figure 2B**). This analysis revealed that there is a module of genes that is upregulated in both cluster 2 and cluster 3, relative to cluster 1, but also that a subset of these genes is dramatically more upregulated in cluster 3 compared to both cluster 1 and cluster 2.

**Figure 2:**
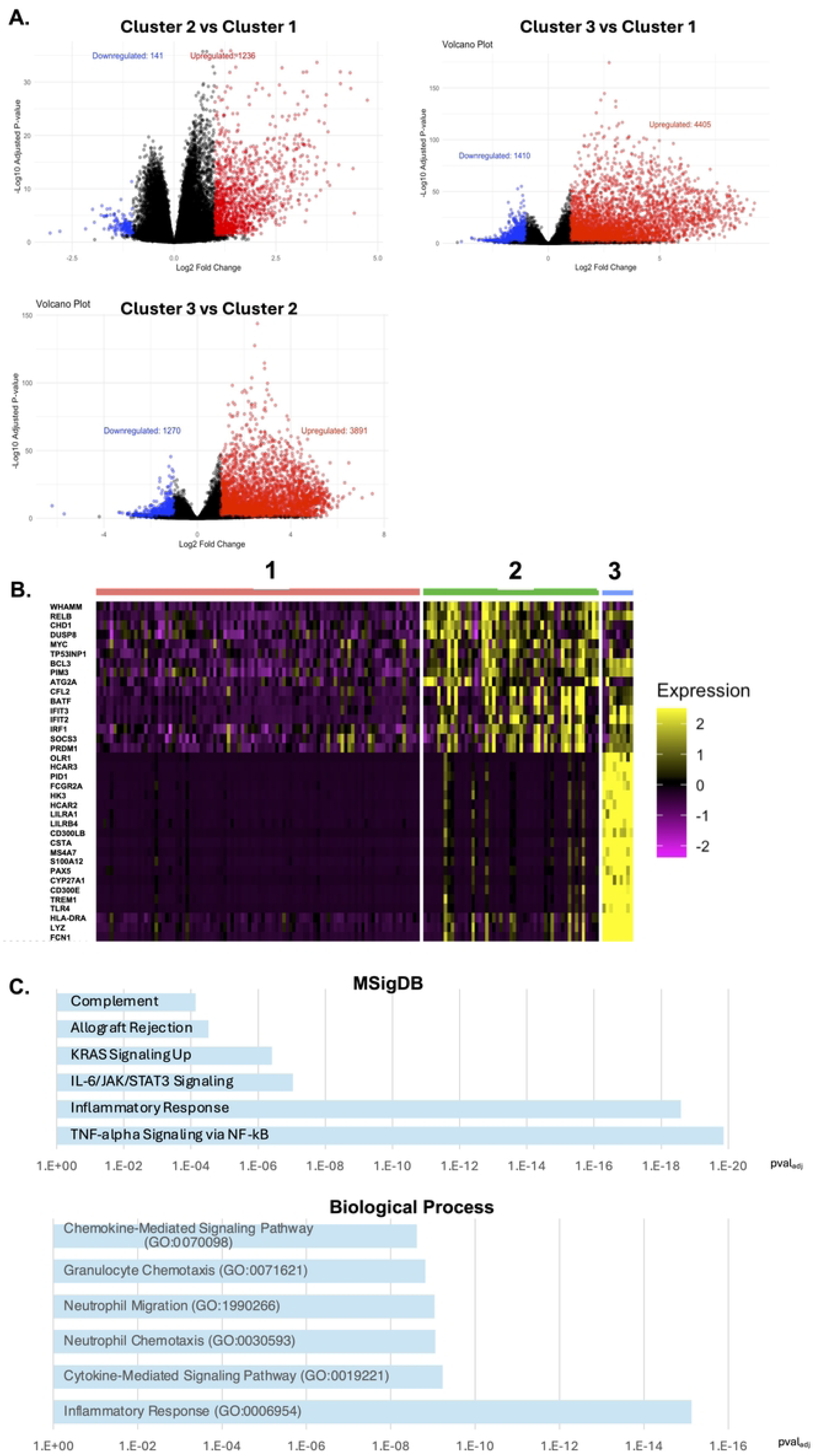
Identification of differentially expressed genes across PWH clusters. **A.** Volcano plots comparing gene expression for different transcriptomic PaCMAP clusters. Upregulated genes (Pval_adj_<0.05, Log_2_FC>1) shown in red, downregulated genes shown in blue. **B.** Heatmap of expression for the top 20 differentially expressed genes across the PaCMAP clusters. Cluster membership shown at top and gene names on the y axis. **C.** MSigDB and biological process enrichment analysis for genes upregulated in cluster 2 vs cluster 1.

We then examined the specific biological pathways that are represented in the set of transcripts that distinguish the clusters from each other. For this comparison we focused primarily on the comparison between cluster 1 and cluster 2 due to the more abundant representation of PWH in these two clusters. To achieve this goal, we used ENRICHR [14], using either the top 200 upregulated transcripts or downregulated transcripts as inputs. When we examined top 200 upregulated genes for cluster 2 versus cluster 1 using the ChEA transcription factor (TF) ChIP-seq reference dataset [15], we observed a strong enrichment of genes that have been shown to bind the NF-κB subunit RELB (Pval_adj_=3.6×10^-5^), suggesting that activity of NF-κB is upregulated in this cluster (**Table S5**). Consistent with this hypothesis, Gene ontology (GO) gene sets associated with NF-κB signaling were also highly enriched in cluster 2 upregulated genes along with inflammatory signaling genes. Among the MSigDB Hallmark sets of pathways [16], ‘TNF-α signaling through NF-κB’ was the most highly ranked pathway, followed by ‘inflammatory response’ (**Figure 2C, Table S6**). For ‘Biological Process’, the upregulated DEGs were enriched for genes that participate in ‘Inflammatory Response’ (GO:0006954) and ‘Immune Marker-Mediated Signaling Pathway’ (GO:0019221) (**Table S7**). We also examined the set of genes that were upregulated in cluster 1 relative to cluster 2 using ENRICHR but we did not observe enrichment of any particular molecular pathway or biological process. When we compared cluster 3 to cluster 2 and cluster 1, we observed enrichment of a similar set of inflammatory pathways in the genes that were upregulated in cluster 3, suggesting that these three clusters represent different levels of activity for the same inflammatory pathway.

To further investigate the association between transcriptomic clustering and the NF-κB pathway, we examined the expression of specific genes associated with this pathway within the dataset. NF-κB is a family of transcription factor complexes whose subunits are encoded by a set of five cellular genes (*RELA*, *RELB*, *cREL*, *NFKB1* and *NFKB2*) [17]. When we examined expression of these genes within the data, we observed that both cluster 2 and cluster 3 exhibited significantly higher expression of all five genes compared to cluster 1 (**Figure 3A**). In general, we also observed a trend for higher expression in cluster 3 compared to cluster 2, although this comparison was only significant (P<0.05, Kruskal-Wallis test) for cREL. Activation of the NF-κB pathway also triggers upregulated expression for genes that encode inhibitory proteins that regulate NF-κB activity as part of a negative feedback loop (*NFKBIA*, *NFKBIB*, *NFKBID*, *NFKBIE*) [18]. When we examined expression of this gene family, we observed a similar pattern to the NF-κB genes – cluster 2 exhibited moderately elevated expression compared to cluster 1, while cluster 3 exhibited more elevated expression compared to both cluster 1 and cluster 2 (**Figure 3B**). We next examined expression of a set of immune marker genes known to be regulated by NF-κB signaling in T cells (*TNFA*, *IL10*, *IL1A*, *IL1B*, *IP10*, *CSF3*, *CCL5*, *IFNG*, *IL2* and *IL7*). With the exception of IL-2, all of these genes exhibited strongly elevated expression in cluster 3 relative to both cluster 1 and cluster 2, while two of them (*IP10* and *IL1B*) were also elevated in cluster 2 relative to cluster 1. Interestingly, *IL2* was expressed at a significantly lower level in cluster 3 compared to both cluster 1 and cluster 2, possibly reflecting a higher level of T cell exhaustion [19]. Overall, these data indicate a pattern of distinct levels of NF-κB activity and expression of NF-κB response genes across the clusters, with cluster 1 exhibiting low activity, cluster 2 exhibiting intermediate activity, and cluster 3 exhibiting high activity.

**Figure 3:**
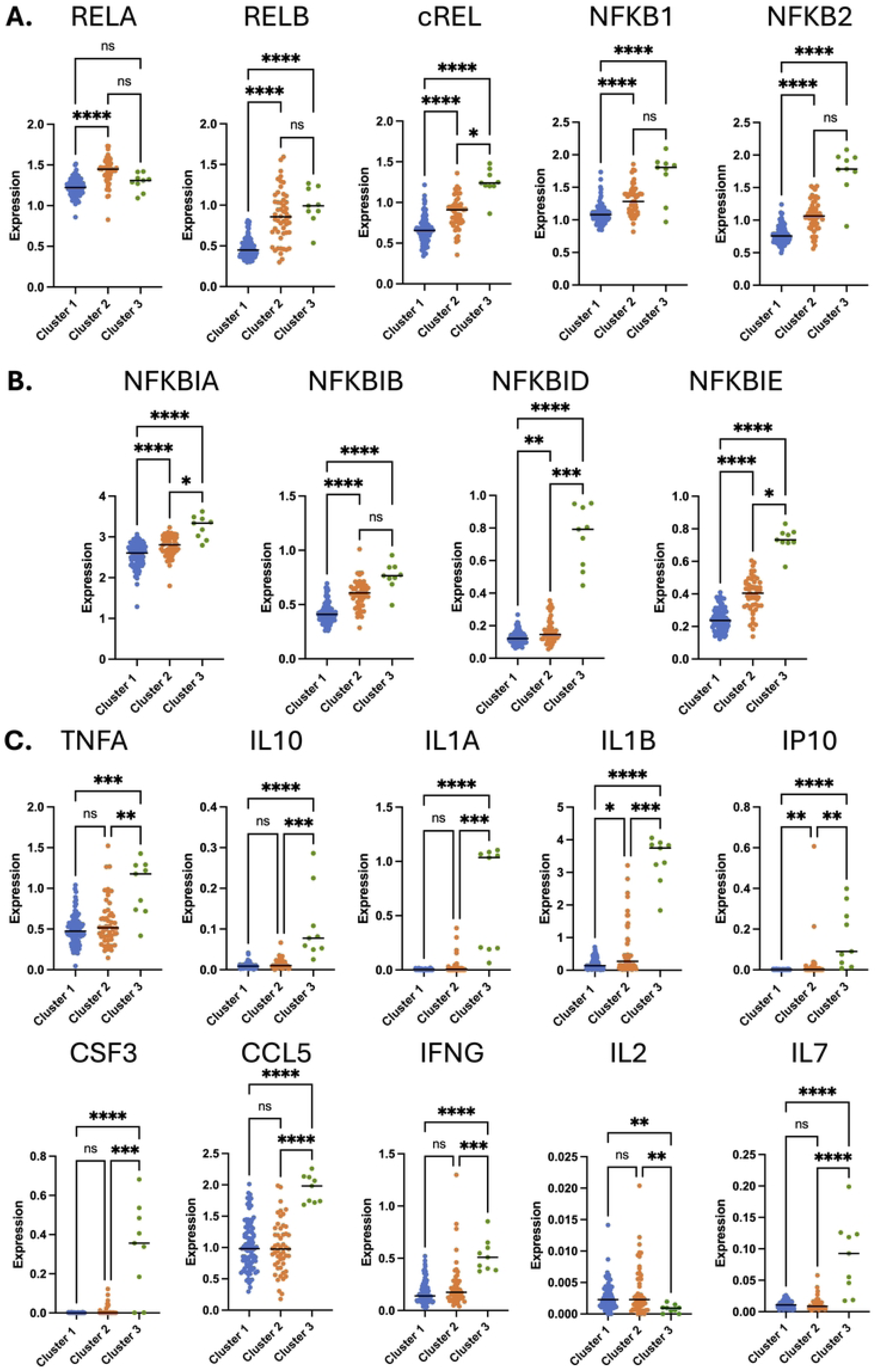
Differential expression of NF-κB regulated genes across transcriptomic clusters. Scaled and natural log normalized gene expression is shown for NF-κB family member genes (**A**), genes encoding NF-κB inhibiting I-κB proteins (**B**), and NF-κB regulated cytokines (**C**) are shown. Each datapoint represents an individual PWH. Cluster 1 – blue, Cluster 2 – orange, Cluster 3 – green. Asterisk indicates statistical significance (P<0.05, Kruskal-Wallis test). ns = not significant (P>0.05). P < 0.05*, P < 0.01**, P < 0.001***, and P<0.0001****.

### Plasma immune marker levels are associated with transcriptomic cluster membership

Since CD4 T cell gene expression is affected by soluble factors such as cytokines in the plasma, we next examined whether the concentration of any of the plasma immune markers measured in this study were associated with transcriptomic cluster membership. Notably, of the seven soluble markers that were measured (IP-10, G-CSF, Pentraxin-3, IL-1β, IL-10, TNF-α, sTLR4), three of these exhibited significantly different levels in the plasma when compared between the clusters (G-CSF, IL-1β, TNF-α), while four were not significantly different (IL-10, IP-10, Pentraxin-3, sTLR4) (**Figure 4**). In particular, IL-1β exhibited a strong difference between clusters, with cluster 2 and 3 having a significantly higher plasma concentration than cluster 1 (median value of 41 fg/mL for cluster 1, 323 fg/mL for cluster 2, 233 fg/mL for cluster 3, P value<0.0001 Kruskal-Wallis test). TNF-α was modestly elevated in cluster 2 versus cluster 1 (median value 232 fg/mL in cluster 2 versus 186 fg/mL in cluster 1, P value = 0.018), but much higher in cluster 3 (median value 10869 fg/mL, P value = 0.0015). By contrast, G-CSF was modestly but significantly higher for members of cluster 1 compared to cluster 2 (8.19 pg/mL for cluster 1, 6.87 pg/mL for cluster 2, P value = 0.0113). Interestingly, the anti-inflammatory cytokine IL-10 trended higher for cluster 1 but this difference was not statistically significant. These data demonstrate that, in addition to the cluster membership deriving from differential expression of inflammation-associated NF-κB - regulated genes, individual clusters are also characterized by distinct patterns of plasma immune marker abundance – particularly for IL-1β, but also for TNF-α and G-CSF. Notably, IL-1β and TNF-α are potent inducers of NF-κB, providing a potential functional connection between these two observations [17]. We also examined the number of cellular genes whose expression in CD4 T cells across the entire cohort was positively correlated with the concentration each plasma immune marker, and observed that, when we used a correlation coefficient threshold of 0.3, TNF-α and IL-1β exhibited the highest number of correlated genes (1608 and 697 respectively), consistent with the hypothesis that these cytokines play a dominant role in driving CD4 T cells gene expression in PWH on ART (**Figure S3A, S3B**). Interestingly, when we used a more stringent correlation threshold (>0.5), IL-10 exhibited the highest number of correlated genes (92 genes) (**Figure S3C**), suggesting that IL-10 regulates a specific module of genes that are closely correlated with this cytokine, but that that TNF-α and IL-1β control a larger set of genes that are more loosely correlated.

**Figure 4:**
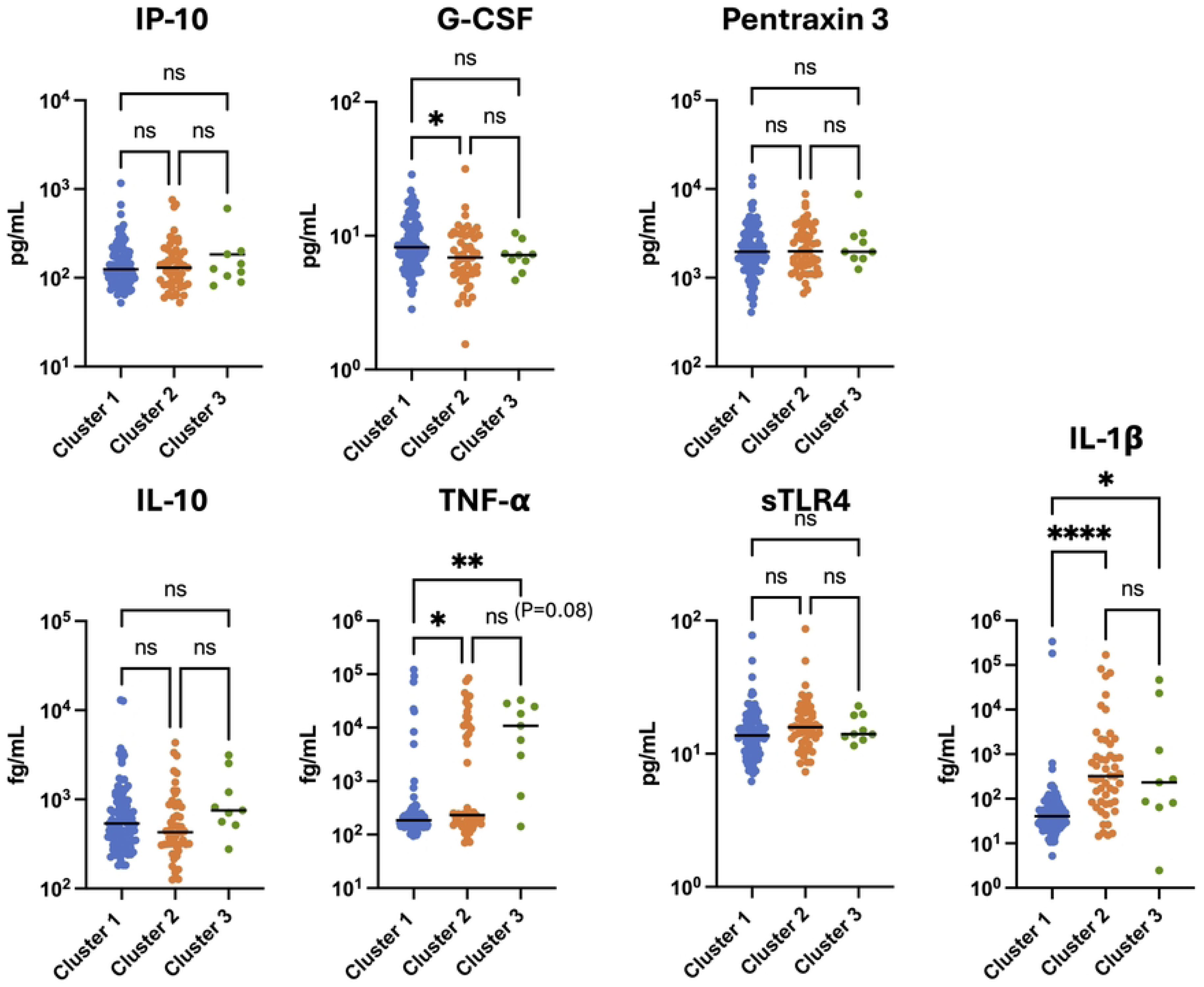
Plasma IL-1β and TNF-α levels are associated with transcriptomic cluster membership. Plasma immune marker concentrations for each member of the cohort are shown, divided by each transcriptomic cluster (Cluster 1 - blue, Cluster 2 - orange, Cluster 3 - green). Each datapoint represents an individual PWH. Asterisk indicates a statistically significant difference between columns (P<0.05, Kruskal-Wallis test). ns = not significant (P>0.05). P < 0.05*, P < 0.01**, P < 0.001*** and P < 0.0001****.

### Cluster membership is not associated with HIV reservoir size

Our dataset of 154 PWH on ART also included HIV reservoir measures: total HIV DNA reservoir size by qPCR, intact HIV DNA by the intact proviral DNA assay (IPDA) [20], and viral RNA (vRNA) expression in CD4 T cells by qPCR, as previously described [11]. We examined whether any HIV reservoir measures were different between members of the three clusters (**Figure 5**). For both the total HIV DNA reservoir frequency and the intact reservoir frequency, there was no significant difference between the clusters. HIV cell-associated vRNA levels trended lower in clusters 2 and 3 although this difference did not reach statistical significance (P=0.12 for cluster 3 vs cluster 1, Kruskal-Wallis test). However, when we combined the clusters with elevated inflammatory gene expression (2 and 3) and compared this in aggregate to cluster 1, we observed that cluster 1 exhibited elevated vRNA compared to the combined 2/3 cluster (P=0.041, Mann-Whitney test) (**Figure S4**). These observations indicate that cluster membership was not associated with the size of the total or intact reservoir but was associated with the transcriptional activity of the reservoir, consistent with previous analyses indicating that HIV RNA levels are inversely correlated with inflammation in PWH on ART [11]. We also observed that the median age of cluster 2 trended higher (median of 49 for cluster 2 vs 45.5 for cluster 1) but this difference was not statistically significant. However, cluster 2 exhibited a lower nadir CD4+ T cell count (median 291 vs 354) and lower median pre-ART viral load (23800/mL vs 57823/mL) than cluster 1. These results indicate that pre-ART clinical parameters of viral load and nadir CD4+ T cell count are associated with membership in cluster 1 vs cluster 2. Cluster 3 was not significantly different from either cluster 1 or cluster 2 for any of the parameters examined, likely due to the smaller number of PWH in this cluster.

**Figure 5:**
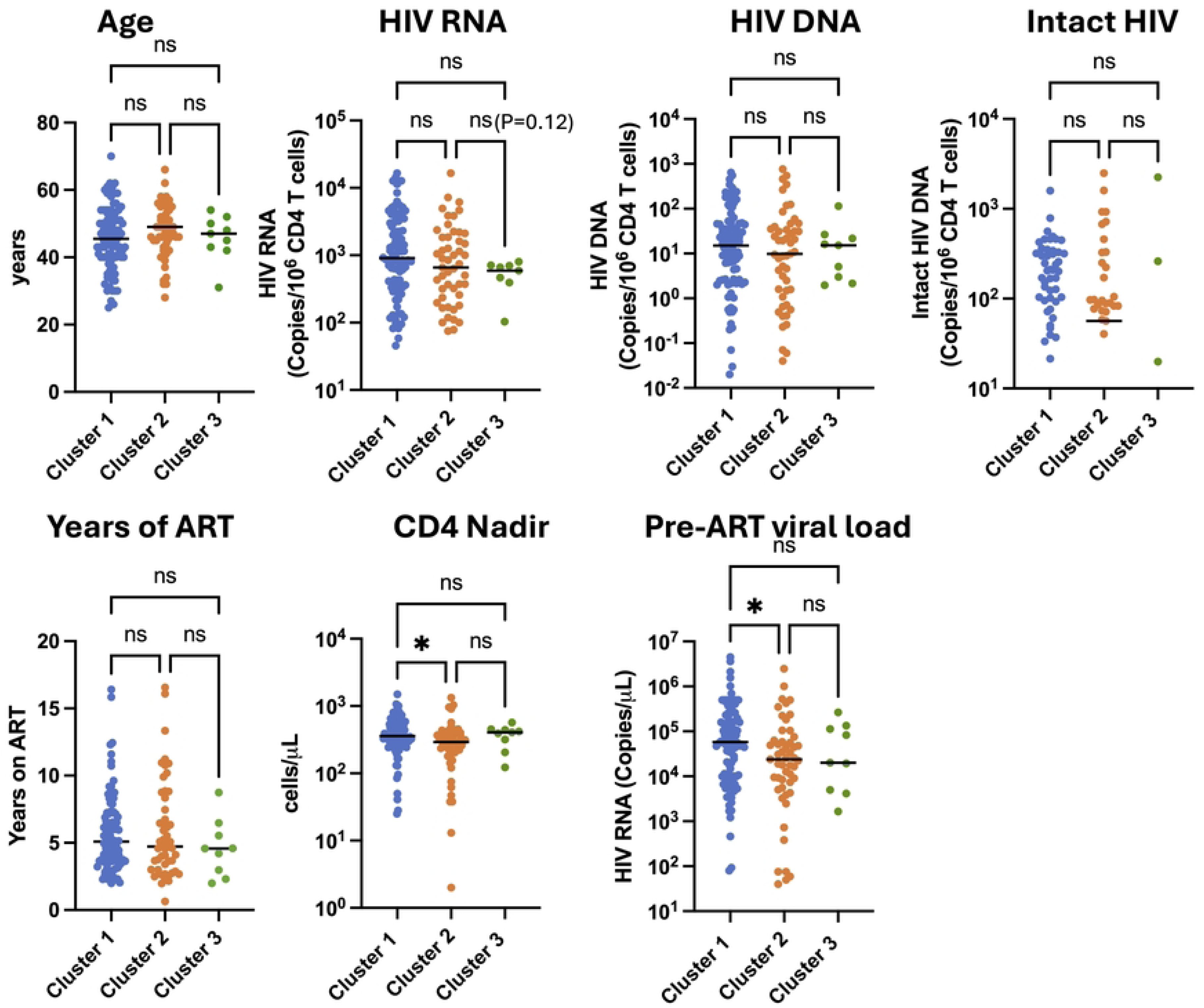
Cluster 2 is associated with a lower pre-ART viral load and nadir CD4+ T cell count. Viral and clinical features for each member of the cohort are shown, divided by each transcriptomic cluster (Cluster 1 - blue, Cluster 2 - orange, Cluster 3 - green). Each datapoint represents an individual PWH. Asterisk indicates a statistically significant difference between columns (P<0.05, Kruskal-Wallis test). ns = not significant (P>0.05).

### Clustering is associated with CD4:CD8 ratio, a marker of host immune recovery and predictor of mortality in PWH on ART

We then examined whether cluster membership was associated with the abundance of any of the major immune subsets present in peripheral blood. For a subset of the cohort (n=66), whole blood was analyzed by routine clinical immunological screening on the day that samples were drawn for both reservoir analysis and RNAseq. This screen measured the abundance of CD4 T cells, CD8 T cells, monocytes, neutrophils, and eosinophils per μL of blood, as well as hemoglobin levels. Since only three members of cluster 3 were represented in this subset, this cluster was not included in further analysis. When we compared cluster 1 to cluster 2, we observed that most immune cell types were not differentially abundant between the clusters (**Figure 6**). However, we observed that cluster 2 exhibited a significantly lower CD4:CD8 T cell ratio compared to cluster 1 (median 0.78 vs 1.04, p=0.0491 Mann-Whitney test). Additionally, members of cluster 2 had an elevated frequency of basophils in their blood compared to cluster 1 (median 0.85% vs 0.6%, p=0.0155 Mann-Whitney test), although this result was largely driven by a single outlying participant. These data thus demonstrate that the CD4 T cell transcriptomic clusters identified by PaCMAP, in addition to their association with plasma immune marker levels, are also associated with specific alterations to the abundance of cell types in the blood of PWH on ART. The reduced CD4:CD8 T cell ratio in cluster 2 could potentially be associated with adverse clinical outcomes for PWH based on cluster membership.

**Figure 6:**
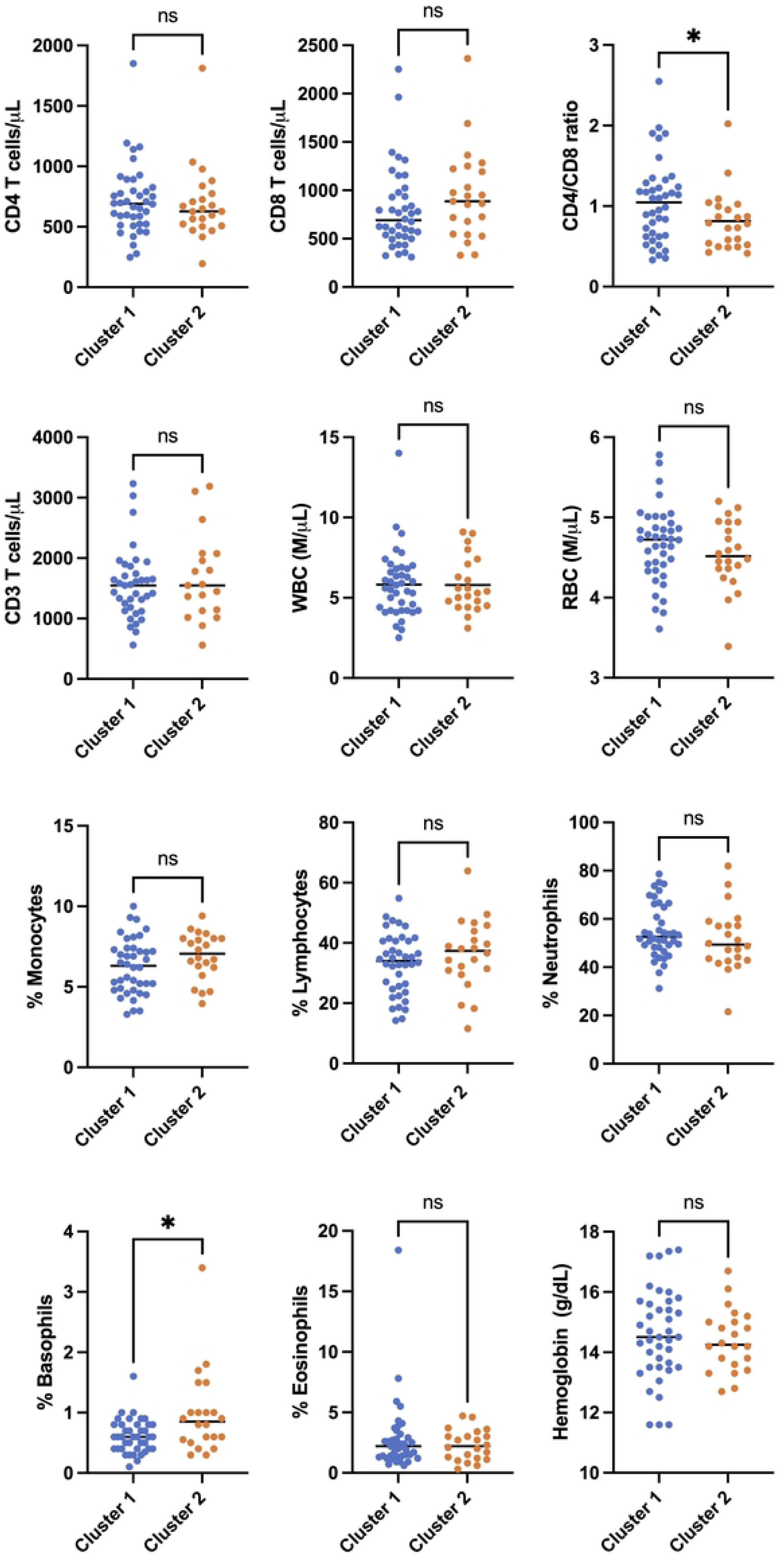
Cluster membership is associated with CD4:CD8 T cell ratio in peripheral blood. Blood cell abundances of selected cell types are shown for a subset of the cohort (n=66) are shown for each transcriptomic cluster (Cluster 1 - blue, Cluster 2 - orange). Each datapoint represents an individual PWH. Asterisk indicates a statistically significant difference between columns (P<0.05, Mann-Whitney test). ns = not significant (P>0.05). P < 0.05*.

### Impact of ART timing on cluster membership

Since our overall cohort consists of a mixture of PWH that were treated with ART during chronic infection and PWH that were treated early during infection (less than 6 months post infection), we next examined whether transcriptomic clustering was associated with the timing of ART initiation, a strong clinical predictor of HIV reservoir size [21]. When we projected the categories of early and late ART timing onto the PaCMAP clustering plot, we observed that early and late treated PWH were found in all three clusters (**Figure 7A**). When we visualized the cluster composition of early vs late treated PWH as a pie chart we observed that, for both early and late treated groups, the majority of participants were found in cluster 1 (**Figure 7B**). Nevertheless, for early treated participants, a lower proportion were found in cluster 2 compared to late treated participants (20% vs 37%), suggesting an impact of ART timing on cluster membership (P=0.0635, Fisher’s exact test). Similarly, when we visualized the data by cluster, the majority of PWH in all three clusters were late treated PWH, reflecting their higher overall abundance in the cohort, but cluster 1 contained a somewhat greater fraction of early treated participants (29% for cluster 1, versus 14% for cluster 2 and 11% for cluster 3, **Figure 7C**). Thus, while timing of ART may have a moderate impact on cluster membership, it does not fully explain the overall clustering pattern.

**Figure 7:**
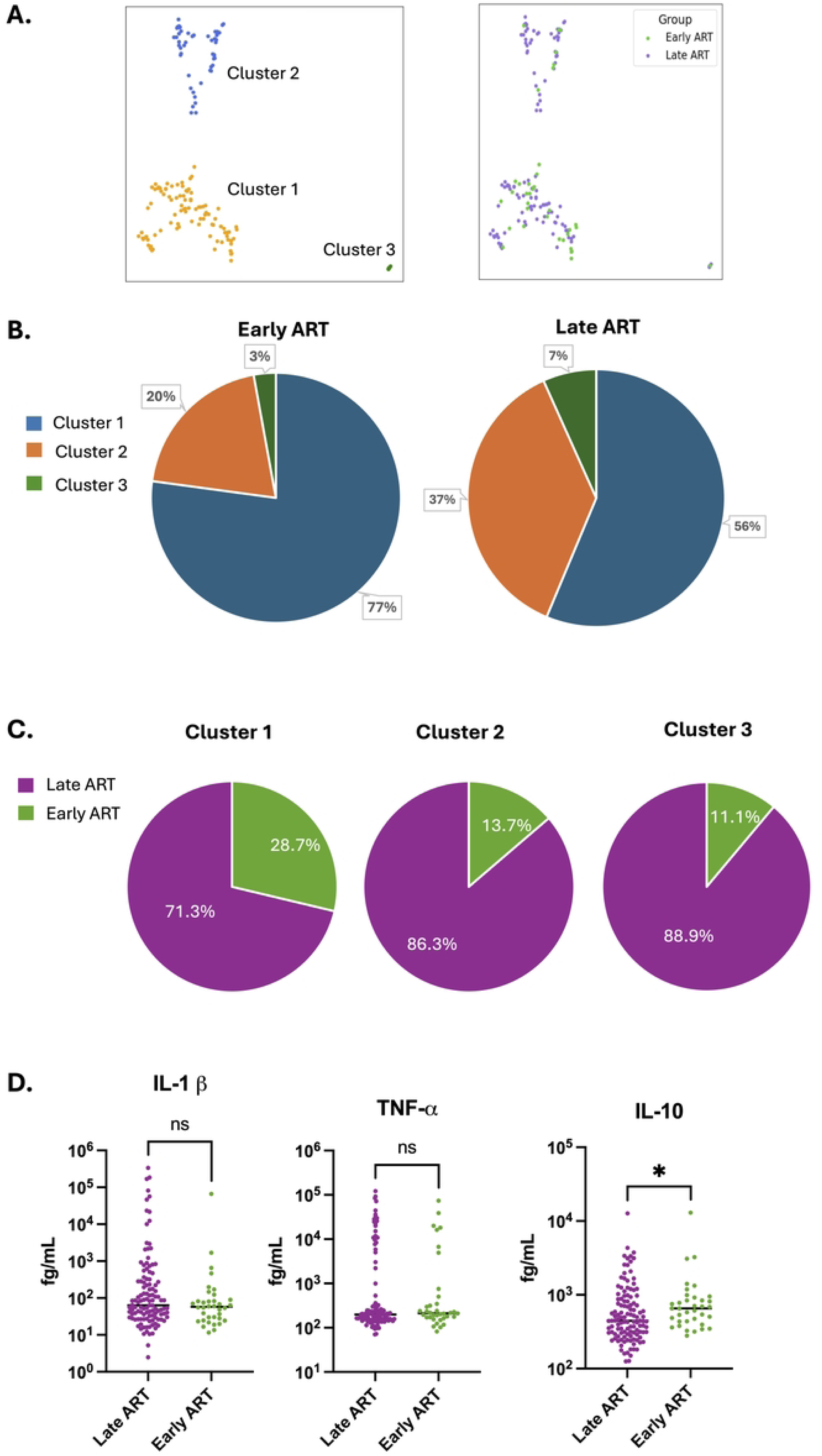
Timing of ART is partially associated with cluster membership but not with plasma IL-1β or TNF-α levels. **A.** PaCMAP projection of the PWH cohort, color-coded by cluster membership (left panel) and by timing of ART initiation (right panel). Early ART treatment (<6 months post infection shown in green, late ART treatment (>6 months post infection) shown in purple. **B.** Pie charts of the cohort, subdivided by timing of ART initiation, showing cluster representation in each group. **C.** Pie charts of the cohort subdivided by cluster membership, showing timing of ART for each group. **D.** Plasma cytokine concentrations for each member of the cohort are shown, divided by timing of ART initiation. Each datapoint represents an individual PWH. Asterisk indicates a statistically significant difference between columns (P<0.05, Kruskal-Wallis test). ns = not significant (P>0.05). P < 0.05*.

We also examined the differential abundance of plasma immune markers between early and late treated PWH. Notably, the abundances of the two inflammatory cytokines (IL-1β and TNF-α) that are the most highly associated with cluster membership were not significantly different between the early and late treated PWH groups (**Figure 7D**). Interestingly, the abundance of the anti-inflammatory cytokine IL-10 was significantly higher in the early treated cohort. Other immune markers that had been measured were not different between the early and late treated groups (**Figure S5**). As previously observed, we found that HIV DNA and RNA were significantly lower for early treated PWH (**Figure S5**) [22–24]. Overall, our data indicate that timing of ART has a moderate impact the transcriptome of CD4 T cells in PWH, and that plasma abundance of the primary transcriptomic cluster-associated cytokines (IL-1β and TNF-α) are independent of the timing of ART.

## Discussion

In this study, we identified three unique clusters of people with HIV on ART using a novel clustering method, bulk RNA sequencing from CD4+ T cells, and plasma immune biomarkers. Using a novel dimension reduction method (PaCMAP), we found three clusters distinguished by a specific biological pathway, NF-κB. Cluster 2 exhibited moderately elevated expression of NF-κB pathway-associated genes, while another cluster (cluster 3) exhibited greatly elevated expression of many of these same genes and also numerous NF-κB regulated cytokines. Furthermore, this phenotype was associated with elevated (eight-fold) plasma IL-1β levels for cluster 2 and 3, while cluster 3 also exhibited greatly upregulated (∼50 fold) plasma levels of TNF-α. Since IL-1β and TNF-α are known to activate NF-κB, these findings suggest that the transcriptomic signature observed in cluster 2 may be driven by the elevated abundance of plasma IL-1β in members of this cluster, while the signature of cluster 3 may be driven by a combination of IL-1β and TNF-α. Thus, we hypothesize that, for a subset of PWH on ART, IL-1β and/or TNF-α expression remain persistently high during therapy, and that the presence of these cytokines in plasma drives chronic expression of an inflammatory signature in the immune cells of PWH via NF-κB. These clusters might then represent distinct inflammatory states – low (cluster 1), intermediate (cluster 2) and high (cluster 3) that are driven by the combination of these immune markers (**Figure 8**). Notably, we observed that members of cluster 2 exhibited a lower CD4:CD8 T cell ratio compared to cluster 1. The CD4:CD8 ratio has been previously shown to be predictive of mortality and non-AIDS morbidity for PWH, even during suppressive ART [8–10,25]. This observation thus suggests that cluster membership may also predict these same comorbidities. It remains unclear whether PWH in cluster 3 also exhibit an altered CD4:CD8 ratio – we were unfortunately unable to compare cluster 3 to clusters 1 and 2 for this parameter due the small number of participants in this cluster with available CD4:CD8 data (N=3). It is possible that analysis of a larger cohort could clarify this question.

**Figure 8:**
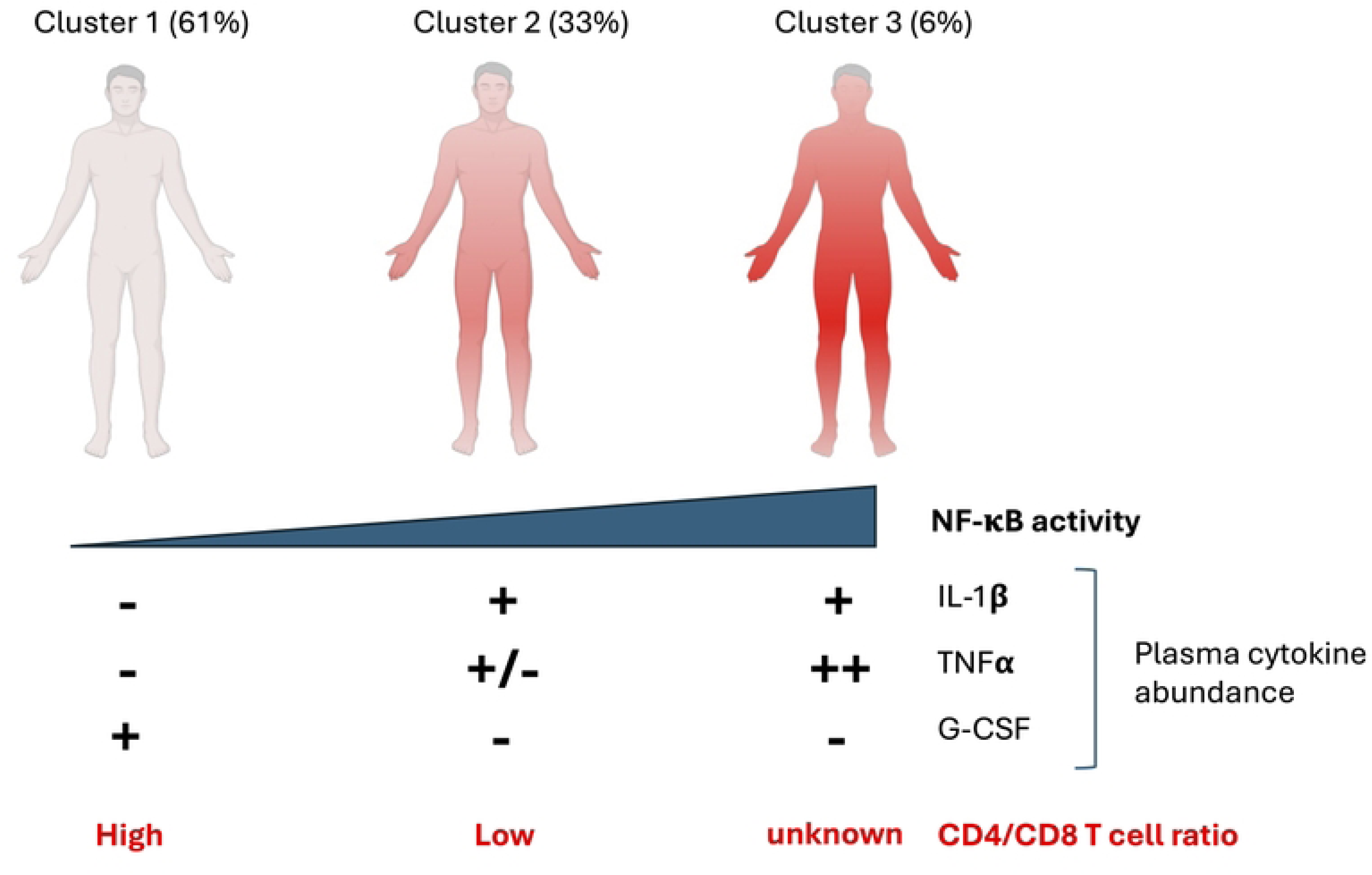
Model for cytokine abundance and CD4 T cell transcriptome clustering. Conceptual model shown for how plasma immune marker abundance and NF-κB activity associates with transcriptomic cluster for CD4 T cells, immune cells and overall inflammation level in people with HIV (PWH). Figure prepared using Biorender.

Furthermore, clinical factors associated with enhanced immune recovery (e.g., lower pre-ART viral load, higher nadir CD4+ T cell count, and earlier timing of ART initiation) [26] were also associated with cluster membership. Thus, it is possible that virological and immunological events prior to therapy establish a pro-inflammatory environment that persists during therapy for a subset of PWH. Notably, we observed no association between the cluster membership and the size of the total DNA reservoir or the intact DNA reservoir. We observed a trend towards lower HIV RNA levels in clusters with elevated expression of NF-κB related genes, consistent with prior observations that viral RNA expression is inversely associated with inflammatory gene expression [11]. This curious association warrants further investigation, since NF-κB is well known as a positive regulator of HIV expression [27,28]. We speculate that elevated NF-κB activity in CD4 T cells may accelerate negative selection of transcriptionally active proviruses over time.

Our study should be considered in the light of several caveats and limitations. The dataset we have analyzed is cross sectional and does not contain any information about longitudinal dynamics for any of the phenomena we observe. The highly elevated clusters such as cluster 3 could, for example, potentially represent a transient state due to a secondary infection. Nevertheless, the transcriptomic clusters identified by PaCMAP were not associated with age or time on therapy, indicating that these phenotypes may be stable during ART. As with any correlational study, it is possible that unmeasured confounders, such as genetics and lifestyle factors such as drug use, contribute to the clustering pattern. Also, the cohort for this study was 96% male and may not represent female PWH. This present study only considers the transcriptomic phenotype of bulk CD4 T cells from peripheral blood, and future studies may need to take a broader and higher resolution view of the immune cell phenotypes using methods such as single cell RNAseq, as well as investigate the immune cells and the HIV reservoir in tissues. As yet, the full clinical significance of the PWH clusters we have identified is unknown, and correlating cluster membership with actual clinical comorbidities (such as vascular events, neurocognitive decline, etc.) will be important to establish the connection between transcriptomic clusters and these outcomes. Although we did not detect an association between cluster membership and measurements of either total or intact reservoir size, it is important to note that only a subset of the cohort exhibited detectable intact proviruses by IPDA. Additional analyses with a larger cohort could clarify the relationship between the intact reservoir and the transcriptomic clusters.

Overall, our study sheds light on the diversity of CD4 T cells phenotypes within a cohort of PWH and points to a significant role for plasma IL-1β/TNF-α and activation of the NF-κB pathway in driving transcriptomic phenotypes for CD4 in PWH on ART. Additionally, this study represents the first identification of a transcriptional signature associated with a reduced CD4:CD8 T cell ratio, and as such, this work may shed light on the molecular basis for how this ratio is associated with non-AIDS comorbidities in PWH. The results also highlight the need to better understand subgroups of PWH that may have divergent immunological properties. The identification of these specific subgroups of PWH and the molecular pathways that define them could open up new approaches to stratifying PWH for risk of comorbidities or clinical responses to HIV cure approaches. Novel biomarkers associated with these clusters, for example, could be examined for predictive power for clinical outcomes or responsiveness to therapeutic vaccines or latency reversing agents. Molecular pathways and transcriptional regulators associated with the clustering could also be considered further for targeting by small molecule inhibitors. Additional work will be required to further discern the biological basis for these groups and their clinical significance

## Acknowledgments

This work was supported by the following grants from the National institutes of Health: NIAID #5-R01AI143381, NIAID #5-UM1AI164567, 1R01AI184122-01, KL2TR002317, University of Washington Center for AIDS Research, 5P30AI027757-37, NIDA R01 DA054994, K23GM112526, UM1 AI126623, R01A141003, KL2TR002317, 108072-50-RGRL and RAVEN: INV-008500. The funders had no role in study design, data collection and analysis, decision to publish, or preparation of the manuscript.

## Methods

### Dataset and cohort

The transcriptome, immune marker and HIV reservoir dataset used for our study was previously published and made publicly available [11]. The data are derived from a cohort of 154 virally suppressed PWH that has been previously described and is composed of PWH selected from two cohorts recruited at UCSF (OPTIONS and SCOPE). This data was combined with unpublished data regarding the timing of ART initiation and clinical lab measurements of major immune cell populations for each participant.

### PaCMAP

PaCMAP was implemented as previously described [29]. Before applying PaCMAP to the RNA-seq data from PWH, the data was preprocessed using the Seurat package in R. The preprocessing involved normalizing the data with the “LogNormalize” method and a scale factor of 10,000. Then, the 2,000 most variable features were selected using the “FindVariableFeatures” function with the “vst” selection method. Afterward, in the Python environment, the data was further scaled using a standard scaler. PaCMAP was then applied, with the number of neighbors (“n_neighbors”=6, reduced from the default of 10) adjusted to account for the small sample size of patients. The resulting PaCMAP embedding was then visualized using the Matplotlib package and the “scatter” function in Python.

### DEG analysis

For the differentially expressed genes (DEG) analysis, a DESeq object was created using the raw counts from PWHs’ RNA-seq data, cluster labels derived from the PaCMAP embedding, and the DESeq2 library in R, utilizing the DESeqDataSetFromMatrix function. By setting the “contrast” parameter to compare each pair of PaCMAP clusters, differentially expressed genes (DEGs) were identified between clusters.

### Statistical analyses

Data were analyzed and visualized using Prism. For most comparisons between clusters, data distribution was assumed to be non-normal, and Mann-Whitney or Kruskal-Wallis tests were performed to determine statistical significance.

## Notes

### Competing Interest Statement

The authors have declared no competing interest.

